# Antibody-mediated feedback modulates interclonal competition in the germinal center

**DOI:** 10.1101/2025.11.06.686519

**Authors:** Alexandru Barbulescu, Jana Bilanovic, Tom Langelaar, A. Katharina Teetz, Linas Urnavicius, Alvaro Hobbs, Jin-Jie Shen, Nadine H. Abrahamse, Renan V. H. de Carvalho, Luka Mesin, Cecilio L. Ferreira, Juliana Bortolatto, Gabriel D. Victora

**Author notes:** These authors contributed equally.

## Abstract

Serum antibodies from prior immune responses regulate B cell activation and germinal center (GC) access upon recall immunization. However, how antibodies produced by an ongoing immune response influence the outcomes of contemporaneous GCs is less clear. To explore this, we developed mouse models enabling the targeted ablation of plasma cells and antibodies produced by an immune response of interest, without affecting those produced homeostatically or by prior antigen encounters. Our findings show that, whereas antibody-mediated feedback is not required for affinity maturation, it can influence competition between B cells with different epitope specificities, specifically by reducing the abundance of clones that recognize the same epitopes as circulating antibodies. This modality of feedback represents a mechanism by which antibody responses can influence epitope specificity in ongoing GCs. These findings may therefore have implications for vaccination strategies aimed at steering clonal selection towards desired epitopes on complex antigens.

## INTRODUCTION

Affinity maturation of antibody responses takes place in germinal centers (GCs), microanatomical structures located within secondary lymphoid organs where B cells undergo somatic hypermutation of their B cell receptors (BCRs) followed by affinity-dependent clonal expansion and differentiation into memory B cells and plasma cells (PCs)^1–4^. Entry of B cells into GCs and their selection within these structures are competitive processes governed by the acquisition of limiting amounts of T cell help^5–8^. Competition among GC B cells can be either *intra*clonal or *inter*clonal. *Intra*clonal competition is well understood—it occurs between members of the same expanded clone (defined here as the expanded progeny of a single naïve B cell bearing a unique V(D)J recombination) and results in B cells that accumulate affinity-enhancing mutations becoming gradually enriched at the expense of lower-affinity variants. Thus, the outcome of intraclonal competition is an increase in affinity of individual B cell clones and their antibodies. On the other hand, the mechanisms that govern *inter*clonal competition—that taking place between two or more clones of cells carrying different V(D)J rearrangements—are much less clear. Immunization with complex multi-epitope antigens recruits tens to hundreds of B cell clones into the same GC structure^9^, making interclonal competition the dominant mode of attrition in the GC. Interclonal competition is therefore a critical determinant of the relative abundance of various antibody specificities in an immune response^1^. However, to date, few if any rules acting specifically on the outcomes of interclonal competition in the GC have been proposed.

Antibodies, the end-product of the GC reaction, play an important role in regulating B cell responses^10,11^. Antibody-mediated feedback has been shown to regulate *inter*clonal competition at the level of B cell activation and GC entry. A number of experiments using transfer of antigen-specific immunoglobulin (Ig) demonstrate that antibodies can both suppress and enhance *de novo* B cell responses, depending on factors such as antigen type and antibody concentration, isotype, and timing of administration^12–15^. For example, access to the GC by adoptively transferred monoclonal B cells is suppressed by prior injection of monoclonal antibodies (mAbs) targeting the same epitope^16,17^ but enhanced by mAbs targeting non-overlapping epitopes^18^. Antibody-mediated feedback was also observed in patients receiving a combination of two therapeutic anti-S protein monoclonal antibodies prior to SARS-CoV-2 mRNA vaccination, as they generate memory responses that shift away from epitopes targeted by the exogenous mAbs, in a striking example of convergence between mouse and human phenotypes^19^. Polyclonal sera elicited by primary immunization can exert similar effects to monoclonal infusion, guiding *de novo* B cell responses to a boost away from epitopes already covered by circulating primary antibodies^20,21^. Altogether, these studies demonstrate that both the specificity and the quantity of pre-existing antibody are important factors in selecting the repertoire of B cell clones that participate in GC responses.

Antibodies have also been proposed to drive intraclonal competition, specifically by setting a threshold that GC B cells must overcome in order to access antigen deposited on follicular dendritic cells (FDCs)^22,23^. According to this model, only B cells harboring BCRs with higher affinity than that of secreted antibodies already bound to the antigen on the FDC surface can capture this antigen and thus survive and proliferate. Indeed, administration of IgM antibodies specific for the hapten 4-hydroxy-3-nitrophenylacetyl (NP) at the onset of the GC reaction can accelerate affinity maturation of the NP response, and mice globally deficient in IgM secretion show impaired affinity maturation of serum antibodies^22,24^. However, the absence of all circulating antibodies also leads to changes in the kinetics of the GC reaction^22^, and thus the relative contributions of the effects of epitope-specific antibody-mediated feedback on the selective pressure acting on GC B cells vs. the more general effects circulating antibodies of any specificity may have on GC magnitude and duration have not been determined. Thus, although it is clear that antibodies can modulate GC B cell responses, further investigation into which exact aspects of intra- and interclonal competition are specifically affected by feedback from antibodies generated by an ongoing B cell response is warranted, especially using loss-of-function models in which only antibodies produced by the response of interest are depleted.

Here, we introduce a series of genetic mouse models designed to investigate the effects of antibody-mediated feedback on intra- and interclonal competition within an ongoing, primary GC response under controlled settings. Using an inducible genetic model that allows targeted depletion of antigen-specific PCs on an otherwise Ig-sufficient background, we show that preventing antibody secretion by a specific B cell clone does not have detectable influence on the rate of affinity maturation of that clone and therefore on the dynamics of intraclonal GC B cell competition. On the other hand, high antibody titers modulate interclonal competition in the GC via epitope masking, favoring B cell specificities that are underrepresented in the serum compartment. Based on these findings, we propose that the antibodies produced by an ongoing immune response may help maintain GC epitope diversity by exerting negative feedback on B cell clones with overlapping specificities.

## RESULTS

### A mouse model for the inducible depletion of antigen-specific plasma cells

To investigate the role of antibodies produced in response to a given antigenic stimulus in the context of a fully polyclonal system, we developed a loss-of-function genetic model that enables us to deplete specifically the antibodies produced by a cohort of B cells currently responding to antigen. To achieve this, we engineered an allele of *Prdm1* (encoding for Blimp-1, the master transcription factor of the PC lineage^25^) in which the coding sequence of the gene was followed by a LoxP-flanked translational stop cassette and then by a porcine teschovirus-1 ribosomal skipping peptide (P2A) and the diphtheria toxin receptor (DTR) (**Figs. 1A** and **S1A**). Upon Cre-mediated recombination, the stop codon is excised and DTR is co-translated with Blimp-1, sensitizing Blimp-1-expressing cells to depletion by diphtheria toxin (DT). When this *Prdm1*^LSL-DTR^ allele was crossed to the B cell-specific Cre driver CD23-Cre^26^, administration of DT resulted in efficient depletion of PCs in both mesenteric lymph nodes (LNs) and bone marrow (**Fig. 1B and S2A**). Following the kinetics of depletion showed that PCs reached a minimum at 2 days post-DT, after which they began to reappear in both mesenteric LNs and bone marrow (**Fig. S2B**). These kinetics were similar for PCs of different isotypes (**Fig. S2C**). Continuous depletion of PCs by daily administration of DT led to gradual loss of circulating IgM and IgG with half-lives (1.2 and 6.9 days, respectively) compatible with known values for these parameters^27^ (**Fig. S2D**).

**Figure 1.**
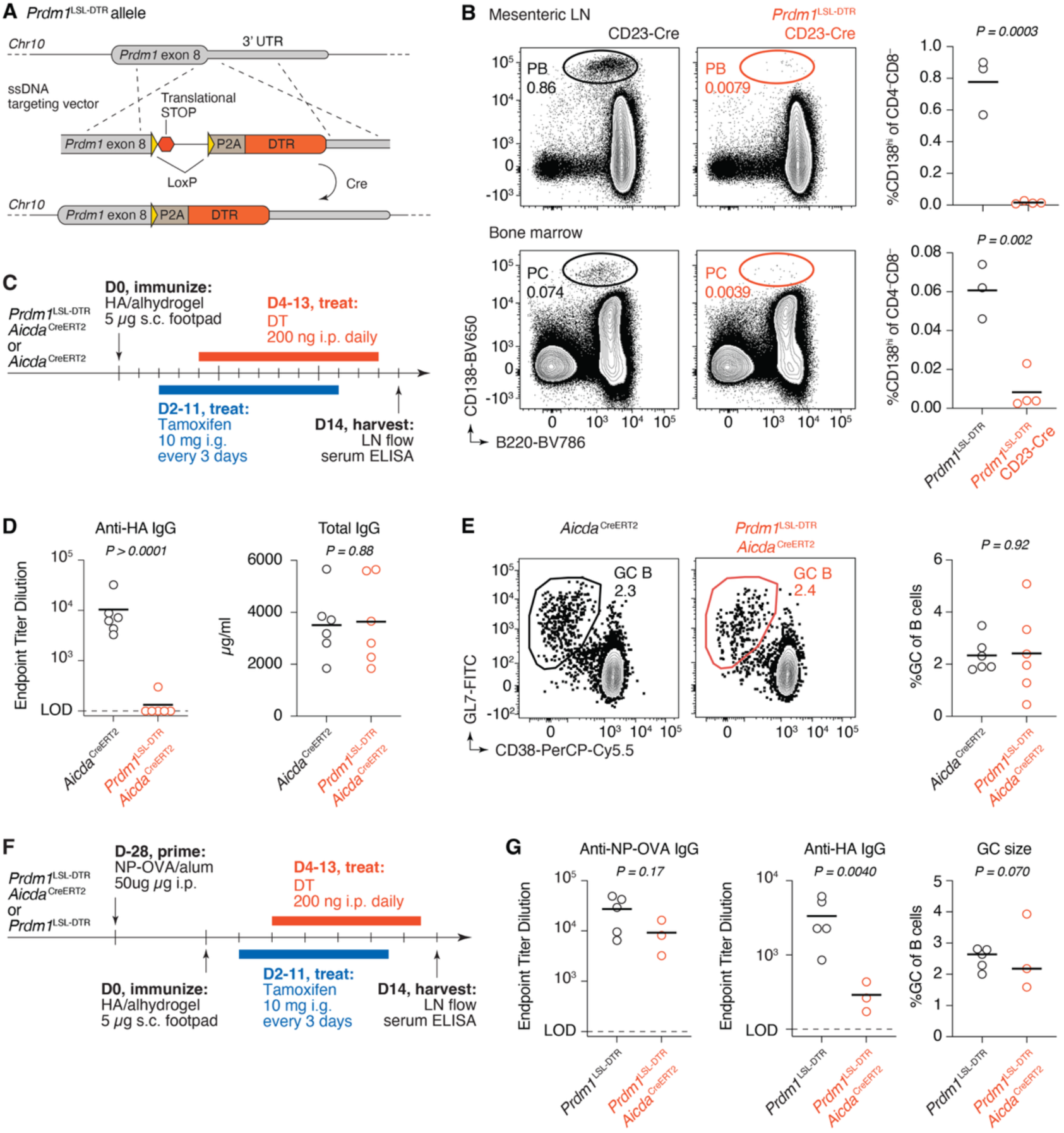
Response specific antibody titers can be depleted using the *Prdm1*^LSL-DTR^ mouse. (A) Schematic of the *Prdm1*^LSL-DTR^ allele. Cre-mediated recombination allows for co-translation of DTR and Blimp-1 from the *Prdm1* locus. **(B)** Depletion of steady-state PCs in mesenteric LNs (*top*) and bone marrow (*bottom*) one day after i.p. administration of 200 ng DT to CD23-Cre.*Prdm1*^LSL-DTR^ mice. **(C)** Experimental setup for panels (D-E). Administration of tamoxifen to *Aicda*^CreERT2^.*Prdm1*^LSL-DTR^ mice recombines the *Prdm1*^LSL-DTR^ allele only in activated B cells, which then become susceptible to killing by DT. **(D)** ELISA quantification of anti-HA IgG endpoint titers (*left*) and total IgG (*right*) at 14 dpi. The limit of detection for endpoint titers was defined at the OD 3 SD above the average OD of blank wells. **(E)** Representative flow cytometry plots of GCs (*left*) and quantification of GC size (*right*) in popliteal LNs 14 dpi. **(F)** Experimental setup for panels (G). **(G)** ELISA quantification of anti-NP-OVA (*left*) and anti-HA (*middle*) IgG endpoint titers, and quantification of GC size (*right*) at 14 dpi. (B,D,E,G) Each symbol represents one mouse. P-values are for Student’s T-test. Results in (D,E) are pooled from two independent experiments, with n = 3 mice each. Results in (G) are pooled from two independent experiments, with n= 1-3 mice per group. Bars in (B, D, E, G) represent means, the bar for the GC size (*right)* in (G) represents median.

To test the ability of the *Prdm1*^LSL-DTR^ allele to specifically ablate serum antibodies elicited by an immunization of interest, we crossed *Prdm1*^LSL-DTR^ mice to the *Aicda*^CreERT2^ strain, in which tamoxifen administration results in Cre-dependent recombination in activated and GC B cells^28^. Treatment of the resulting *Aicda*^CreERT2^.*Prdm1*^LSL-DTR^ mice with tamoxifen removes the stop cassette in ongoing GCs and other activated B cell populations. Plasmablasts and PCs descended from these B cells express DTR and are therefore susceptible to DT-mediated ablation, whereas those derived from steady-state or previous immune responses are preserved. We immunized *Aicda*^CreERT2^.*Prdm1*^LSL-DTR^ mice with recombinant hemagglutinin (HA) from influenza virus strain A/Puerto Rico/8/1934 (PR8) in alhydrogel adjuvant and administered tamoxifen by oral gavage on 2, 5, 8, and 11 days post-immunization (dpi) to ensure maximal recombination in responding B cells. Concurrently, we administered daily doses of DT beginning at 4 dpi (**Figs. 1C**). At day 14, anti-HA IgG titers from most mice were depleted beyond the limit of detection of ELISA, while total serum IgG concentrations remained stable, as PCs produced outside of the HA response were not depleted (**Fig. 1D**). Depletion of plasmablasts in the draining LN required the administration of both tamoxifen and DT (**Fig. S2E,F**). On the other hand, GC size was not affected in the absence of antibody (**Fig. 1E**), indicating that any potential depletion of the small contingent of *Prdm1*-expressing GC B cells^29,30^ by DT is insufficient to detectably alter GC kinetics. Lastly, to confirm that we could use DT administration to specifically deplete an antibody response of interest without affecting a previously established one, we immunized *Aicda*^CreERT2^.*Prdm1*^LSL-DTR^ mice or *Aicda*^CreERT2^ controls with NP coupled to ovalbumin (NP-OVA), immunized mice with HA one month later, and used sequential administration of tamoxifen and DT as above to specifically deplete serum anti-HA antibodies (**Fig. 1F**). This resulted in specific depletion of the anti-HA response, while antibodies to NP-OVA were modestly but not significantly affected (as expected if residual GC responses to NP-OVA are still ongoing at the time of tamoxifen administration). There was again no change in GC size between treatment groups (**Fig. 1G**). We conclude that *Prdm1*^LSL-DTR^ mice provide a tool to specifically and acutely deplete plasmablasts and PCs generated by an immune response of interest, without affecting global Ig concentrations or antibodies derived from prior immune responses. However, given the long half-life of IgGs, ablation of antigen-specific antibody is most efficient when DT is administered prior to the establishment of a robust serum response.

### Antibody-mediated feedback is not required for efficient affinity maturation

To determine the impact of antibody-mediated feedback on intraclonal competition, we used immunization with NP-OVA, which generates GC responses in which affinity can be inferred from *Ig* sequence. Responses to NP-OVA are dominated by B cells carrying the VH1-72 heavy chain V-segment paired to an Igλ light chain. These “B1-8-like” B cells affinity mature by accumulating a stereotypical tryptophan to leucine mutation in CDR1 (W33L) that confers a roughly 10-fold gain in affinity to NP^31^, as well as K59R, a second mutation known to enhance affinity for NP^32^. We immunized *Aicda*^CreERT2^.*Prdm1*^LSL-DTR^ mice with NP-OVA in the footpad and followed the same tamoxifen and DT administration protocol as with HA immunization (**Figs. 2A**). At day 14 post immunization, serum antibody reactivity to NP-OVA and NP-bovine serum albumin (BSA) were nearly entirely depleted in mice that received DT, whereas total serum IgG concentrations remained the same (**Fig 2B,C**). Notwithstanding, there were no significant effects on the proportion of B cells carrying the known affinity-enhancing mutations (**Fig. 2D**).

**Figure 2.**
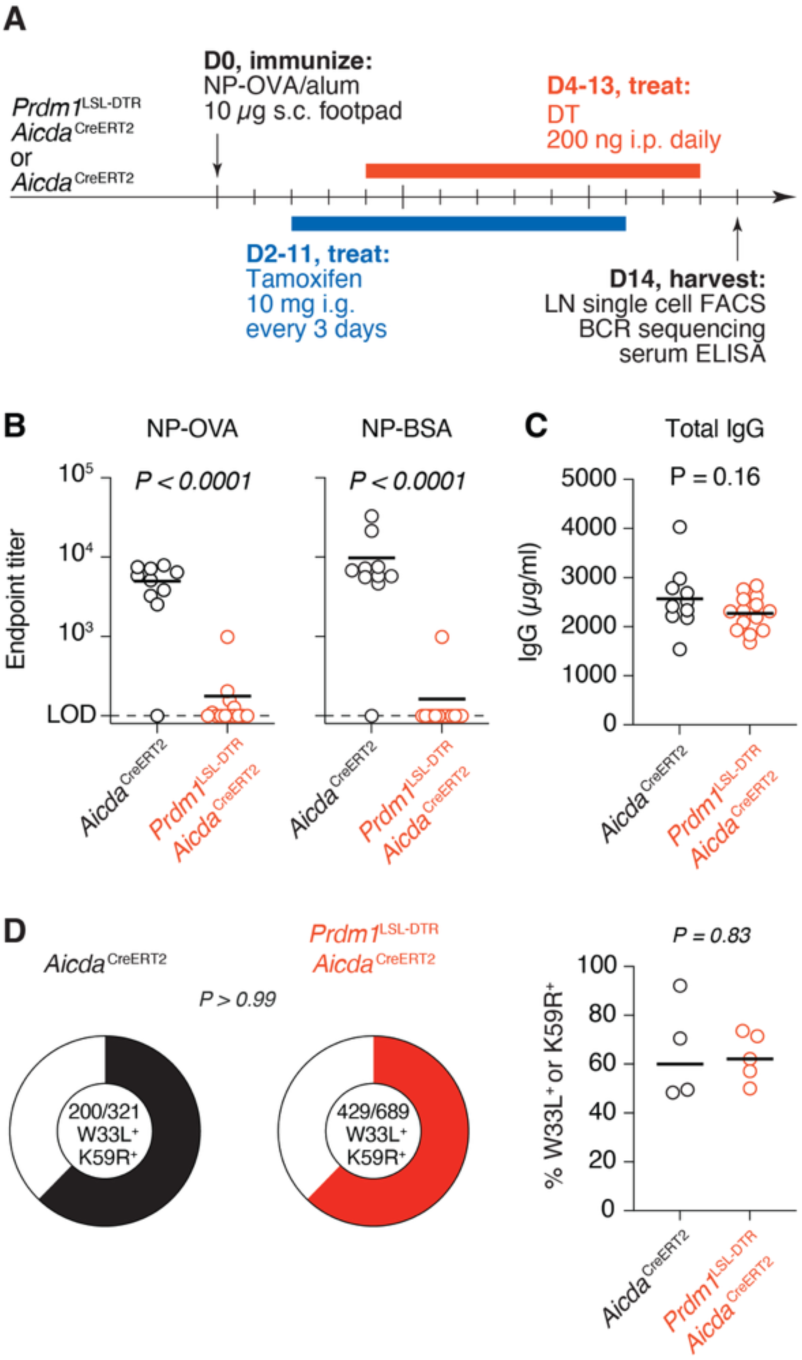
Antibody-mediated feedback does not accelerate affinity maturation in response to the hapten NP. **(A)** Experimental setup. **(B)** ELISA quantification of anti-NP-OVA (*left*) and anti-NP-BSA (*right*) IgG endpoint titers at 14 dpi. **(C)** ELISA quantification of total IgG at 14 dpi. **(D)** *Left*, Pie charts showing proportion of total Vh1-72 B cells sequenced containing affinity-enhancing mutations W33L or K59R. *Right*, percentage of Vh1-72 B cells per mouse sequenced carrying W33L or K59R mutations. (B,C) Data are pooled from two independent experiments, with n = 5-7 mice per group. Bars represent means. (D) Results are pooled from two independent experiments, with n = 2-3 mice per group. Bars represent medians. Each symbol represents one mouse. P-values are for Student’s T-test (columns) and Fisher’s exact test (pie charts). Sequences are available in Supplemental Spreadsheet 1.

To extend these findings to a relevant protein antigen, we generated mice carrying pre-rearranged *Igh* and *Igk* V-regions specific for PR8 HA. To identify a suitable HA-binding antibody, we infected C57BL/6 mice with 33 PFU of mouse-adapted PR8 influenza virus and sorted single HA tetramer-binding mediastinal GC B cells at 21- and 45-days post-infection (**Fig. S3A**). We sequenced the *Igh* and *Igk* loci of each cell and selected 20 expanded clones; from each, we synthesized a recombinant monoclonal antibody (mAb) corresponding to the most abundant member of the clone. All but one antibody showed detectable binding to HA by ELISA (**Fig. S3B**). HA contains five major antigenic sites, each commonly targeted in polyclonal B cell responses^33^. To define the epitope specificity of each clone, we tested their binding to panel of five PR8 HA mutants each retaining only one of the five antigenic sites^33^. Several clones bound predominantly to one of the HA variants, and we selected clone 1.1 for further characterization (**Fig. S3C**). mAb 1.1 binds to the “Sa” antigenic site on the HA head, which is immunodominant in the context of HA protein immunization^33^. The Ig of clone 1.1 (hereafter, “clone Sa”) consists of a heavy chain rearrangement of *Ighv1-50*, *Ighd2-3,* and *Ighj4* coupled with an Igκ light chain rearrangement of *Igkv15-103* and *Igkj1*. Clone Sa carries 6 and 1 non-silent mutations in *Igh* and *Igk*, respectively. Using CRISPR-Cas9 zygote micro-injections^34^, we replaced endogenous mouse *Ighj* and *Igkj* loci with prearranged VDJH and VJκ sequences, respectively (**Figs. 3A)**. We crossed the resulting mouse to an *Igk*^−^ strain to decrease rearrangements of the endogenous *Igk* allele and to the CD45.1 allelic marker, thus obtaining the *Igh*^Sa/+^.*Igk*^Sa/–^ CD45.1/1 (*Ig*^Sa^) mice that we used as donors for adoptive transfer experiments. The large majority of circulating blood B cells in *Ig*^Sa^ mice bound HA (**Fig. 3A**).

**Figure 3.**
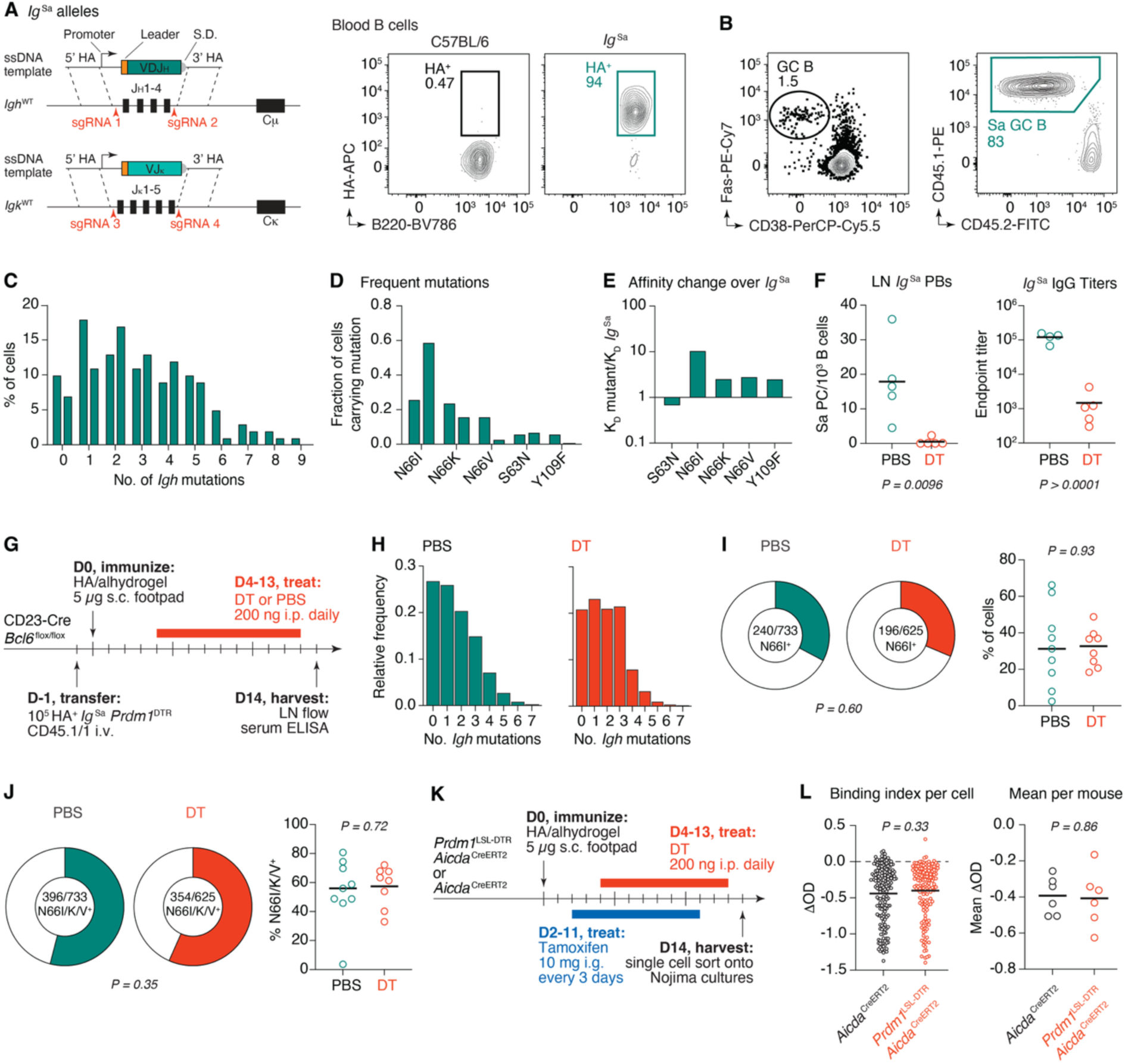
Antibody-mediated feedback does not accelerate affinity maturation in response to influenza HA. **(A)** Generation of the *Ig*^Sa^ BCR knock-in mouse (*left*). Representative flow cytometry plot showing binding to HA tetramers among peripheral blood B cells from an *Ig*^Sa^ mouse (*right*). **(B)** Representative flow cytometry plots showing GC formation (*left*) and presence of transferred B cells in GCs (*right*) in C57BL/6 mice immunized with PR8 HA one day following adoptive transfer of 10^5^ HA^+^ *Ig*^Sa^ B cells. **(C)** Histograms from two mice showing number of *Igh* mutations per *Ig*^Sa^ B cell at 18 dpi. **(D)** Proportion of the five most frequent *Ig*^Sa^ *Igh* mutations in both mice from (C). **(E)** *Ig*^Sa^ mutant Fabs were produced and their affinities to HA measured by bio-layer interferometry. Shown are affinity fold- differences between each mutant Fab and the unmutated *Ig*^Sa^ Fab. **(F)** Flow cytometry quantification of *Ig*^Sa^.*Prdm1*^LSL- DTR^ PCs in popliteal LNs at 7 dpi (*left*) and ELISA quantification of endpoint titers of anti-Sa IgG at 18 dpi (*right*) in the presence or absence of DT treatment. Bars represent means. Data are pooled from two independent experiments, with n = 2-3 mice per group. Each symbol represents one mouse. **(G)** Experimental setup for panels (H-J). **(H)** Distribution of number of *Igh* mutations in *Ig*^Sa^ B cells at 14 dpi in the presence or absence of DT treatment. **(I)** Pie charts show proportion of total cells sequenced containing the affinity enhancing N66I mutation in *Igh* (*left*). Graph shows the percentage of cells per mouse sequenced that harbor N66I (*right*). Each symbol represents one mouse. **(J)** As in I) but including all affinity-enhancing mutations at position N66. Sequences are available in Supplemental Spreadsheet 1. **(K)** Experimental setup for panel (L). **(L)** Binding indices of antibodies produced from single GC B cells. Binding was compared to a that of a high-affinity anti-HA mAb standard (*left*). Each symbol represents one culture. The mean binding index for each mouse is also shown (*right*). Data are pooled from two independent experiments, with n = 3 mice per group. Each symbol represents one mouse. P-values are for Student’s T-test (columns) and Fisher’s exact test (pie charts). (H-J) Results are pooled from two independent experiments, with n = 4-5 mice per group. Bars represent medians.

To characterize the response of *Ig*^Sa^ B cells to immunization, we adoptively transferred 10^5^ HA- binding B cells from *Ig*^Sa^ donors into C57BL/6 hosts one day before footpad immunization with recombinant PR8 HA protein in alhydrogel. *Ig*^Sa^ B cells efficiently entered GCs, accounting for the majority of GC B cells one week after immunization (**Fig. 3B**). Next, we assessed the potential for *Ig*^Sa^ B cells to affinity mature by sequencing the *Igh* loci of transferred B cells 18 days following HA footpad immunization. *Ig*^Sa^ B cells acquired an average of 2.9 *Igh* mutations in the two mice we sequenced, with 26% and 18% of cells containing 5 or more mutations (**Figs. 3C** and **S3D**). Three of the most frequent *Igh* mutations involved the asparagine at position 66 (N66I, N66K, and N66V, present in 42%, 20%, and 10% of *Ig*^Sa^ GC B cells, respectively), suggestive of potential antigen-driven selection for the replacement of asparagine at that position (**Fig. 3D**). Indeed, recombinant fragment antigen-binding regions (Fabs) carrying the N66K, N66V, and N66I mutations showed 2.6, 2.9, and 10.7-fold increases in affinity, respectively, compared to the version of clone Sa carried by *Ig*^Sa^ mice. As expected^35^, the increase in affinity of N66I was driven predominantly by a decrease in the off-rate of the interaction with HA (**Figs. 3E** and **S3E**).

To test the impact of antibody-mediated feedback on affinity maturation in this system, we generated a “pre-floxed” version of the *Prdm1*^LSL-DTR^ allele (*Prdm1*^DTR^) by crossing with *Cd79a*^Cre^ ^36^ and screening for rare offspring in which this driver was active in the germline. We crossed *Prdm1*^DTR^ to *Ig*^Sa^ mice and transferred the resulting B cells into C57BL/6 hosts prior to immunization with HA PR8. We then administered DT or PBS to immunized mice starting at 4 dpi. DT administration markedly depleted *Ig*^Sa^ LN plasmablasts at 7 dpi and, at 18 dpi, Sa-specific titers were approximately 100-fold reduced in DT treated compared control mice (**Fig. 3F**). We next transferred 10^5^ *Prdm1*^DTR^ *Ig*^Sa^ HA-binding B cells into CD23- Cre.*Bcl6*^flox/flox^ hosts^26,37^ (which avoid confounding by endogenous GC B cells) and quantified the appearance of affinity-enhancing *Igh* mutations upon HA PR8 immunization in the presence or absence of anti-Sa antibody titers (**Fig. 3G**). The total number of *Igh* mutations ranged from 0 to 7 at 14 dpi, with no difference in mean values between DT and PBS groups (**Fig. 3H**). Depletion of Sa antibody titers had no detectable effect on the accumulation of N66I or other N66 affinity-enhancing mutations in the GC (**Fig. 3I,J**).

Lastly, we sought to determine the impact of antibody-mediated feedback on the affinity maturation of a polyclonal response to HA. We immunized *Aicda*^CreERT2^.*Prdm1*^LSL-DTR^ mice with HA PR8, administered tamoxifen on 2, 5, 8, and 11 dpi, and concurrently administered DT daily, beginning at 4 dpi (**Fig. 3K**). On day 14, we sorted single HA^+^ GC B cells into “Nojima” cultures^38^ to estimate the avidity of the antibodies these cells encode in the presence or absence of competing serum antibody. Following the *ex vivo* differentiation of the sorted GC B cells into antibody-secreting cells, we harvested culture supernatants, determined the concentration of IgG in each well, and assayed the binding of antibodies in IgG^+^ wells to recombinant HA by ELISA. To compare affinities between B cell clones, we calculated the difference between each experimentally measured optical density (OD) value and the OD obtained for a monoclonal antibody specific for the Cb epitope of HA at the corresponding IgG concentration, interpolated from a standard dilution curve (**Fig. S3F**). The resulting binding indices showed no detectable difference in the average binding to HA of GC B cells generated in the presence or absence of DT treatment (**Fig. 3L**). Thus, experiments using three separate models of affinity maturation failed to detect an appreciable role for circulating antibody in driving affinity maturation in the GC.

### Secreted antibodies limit the competitiveness of GC B cells with the same epitope specificity

A second potential role for antibody-mediated feedback is to regulate competition between GC B cells that bind different epitopes. In such a scenario, antibodies binding to a particular epitope would decrease the effective concentration of that epitope available to GC B cells, providing a competitive advantage to clones specific for non-overlapping sites on the same antigen^39^. We sought to determine whether antibody-mediated feedback could have an effect on interclonal competition along these lines. We first reanalyzed our data from immunization with NP-OVA (**Fig. 2**), in which a large proportion of the GC response is directed toward the immunodominant NP hapten^40^. Ablation of antibody, which was predominantly NP-specific (**Fig. 2B**), resulted in a significant increase in the number of NP-binding GC B cells (measured as the percentage of Igλ^+^ cells, which account for the majority of NP-binding B cells in C57BL/6 mice^41^) 14 days following immunization, while GC sizes remained stable (**Fig. 4A**). This observation suggests that antibody may indeed affect the balance of epitopes targeted by GC B cells.

**Figure 4.**
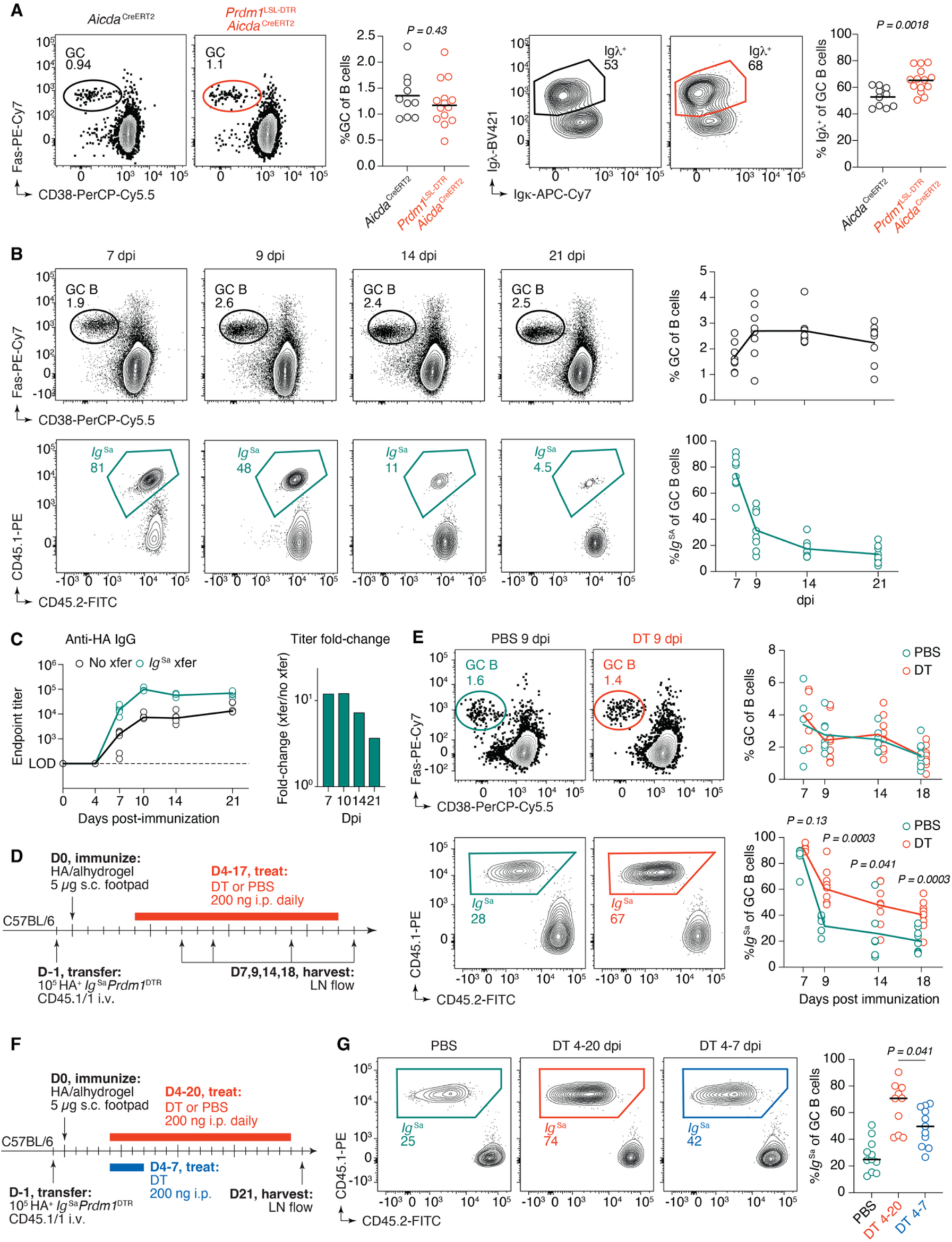
Sa-specific antibody limits the success of clone *Ig*^Sa^ in the GC. (A) Representative flow cytometry plots and quantification of GC size (*left*) and fraction of Igλ^+^ GC B cells (*right*) at 14 dpi with NP-OVA. (B) Immunization time course with HA following adoptive transfer of *Ig*^Sa^ HA^+^ B cells to C57BL/6 mice. Top row shows representative flow cytometry plots and quantification of GC size and bottom row shows representative flow cytometry plots and quantification of *Ig*^Sa^ GC occupancy. (C) Anti-HA IgG endpoint dilution titers following immunization of C57BL/6 mice with HA with and without transfer of *Ig*^Sa^ (*left*). Fold-change in anti-HA IgG endpoint dilution titers in recipients of *Ig*^Sa^ B cells compared to mice without transfer (*right*). Data are pooled from two independent experiments, with n = 3 mice per group. Lines connect medians. **(D)** Experimental setup for panels (E,F). **(E)** Representative flow cytometry plots of GC size at day 9 (*top, left*) and quantification of GC size over time (*top, right*) with and without DT administration. Representative flow cytometry plots of *Ig*^Sa^ GC occupancy at day 9 (*bottom, left*) and quantification of *Ig*^Sa^ GC occupancy over time (*bottom, right*) in the presence or absence of DT treatment. **(F)** Experimental setup for panel (G). **(G)** Representative flow cytometry plots of GC occupancy at day 21 following no depletion of antibody, continuous depletion of antibody, and depletion of only early extrafollicular antibody (*left*) and quantification (*right*). In all column plots, each symbol represents one mouse. P-values are for Student’s T-test. (A) Results are pooled from two independent experiments, with n = 5-7 mice per group. Bars represent means. (B, E) Results are pooled from 3 independent experiments, with n = 2-3 mice per group. Lines connect means. (G) Results are pooled from 2 independent experiments, with n = 5-6 mice per group. Bars represent medians.

To test this in a controlled setting, we performed experiments in which *Ig*^Sa^ B cells were adoptively transferred into C57BL/6 mice. We hypothesized that if antibody-mediated feedback affects the competitiveness of B cells with the same specificity, the *Ig*^Sa^ B cell clone would become progressively less represented in GCs over time due to increasing feedback inhibition by its own antibody. We performed a time course of *Ig*^Sa^ in the GC following transfer into C57BL/6 mice and footpad immunization with HA.

Although GC size remained relatively stable over three weeks, the proportion of transferred cells within GCs progressively contracted (**Fig. 4B**). This decay was already substantial as early as 9 dpi, when *Ig*^Sa^ B cells accounted for on average 32% of the GC, down from over 80% at 7 dpi. By 21 dpi, clone *Ig*^Sa^ represented only 12% of GC B cells (**Fig. 4B**). Similar depletion of specific cells over time was observed when tracking Igλ^+^ cells in GCs responding to NP-OVA, confirming that this suppression is not unique to adoptive transfer systems (**Fig. S4A**).

Quantification of epitope-specific titers following immunization with HA showed that Sa reactivity was on average 9-fold higher when *Ig*^Sa^ B cells are transferred compared to non-transfer settings (**Fig. 4C**). Additionally, nearly all anti-HA IgG reactivity in serum in this model was derived from transferred cells, as Sa-specific titers were almost identical to the total anti-HA titers (**Fig. S4B**). This indicates that the adoptively transferred population outcompetes endogenous B cells for differentiation into antibody secreting cells, as well as for entry into the GC. Increased Sa titers with *Ig*^Sa^ B cell transfer along with the decay of *Ig*^Sa^ in the GC raised the possibility that anti-Sa antibody feeds back onto the GC response, thereby occluding the Sa epitope on intact HA and reducing the competitiveness of B cells that bind that epitope, possibly representing a mechanism to promote epitope diversity within an ongoing GC reaction.

To test this, we repeated the adoptive transfer time-course experiment in **Fig. 4B** but using *Prdm1*^DTR^ *Ig*^Sa^ B cells and depleting antibody produced by the transferred cells beginning at 4 dpi (**Fig. 4D**). Depletion of *Ig*^Sa^-derived antibody throughout the GC response resulted in partial rescue of *Ig*^Sa^ GC B cells, without altering the mean GC size (**Fig. 4E**). Whereas removal of antibody had no effect on GC entry by *Ig*^Sa^ B cells at 7 dpi, the proportion of transferred cells increased from 32%, 26%, and 10% of the GC in controls to 61%, 48%, and 41% in DT-treated mice at 9, 14, and 18 dpi, respectively. This enrichment was not due to non-specific adjuvant effects of DT, as administration of DT during an identical experiment in which transferred *Ig*^Sa^ B cells were *Prdm1*^WT/WT^ did not yield any differences in *Ig*^Sa^ content in the GC at day 18 (**Fig. S4C**). Repeating this experiment using an unrelated BCR knock-in model specific for the model antigen chicken IgY (*Ig^chIgY^*)^42^ showed similar sensitivity to antibody feedback, albeit with delayed kinetics (**Fig. S4D,E**), likely related to the delayed appearance of anti-IgY antibodies in this system compared to *Ig*^Sa^ B cell transfer followed by HA immunization (**Fig. S4F**).

To untangle the effects of early extrafollicular and later GC-derived antibody on feedback inhibition, we repeated the *Ig*^Sa^ B cell transfer experiment but depleted antibodies only from 4 through 7 dpi, blunting the extrafollicular response but sparing GC-derived antibodies produced by plasmablasts generated after day 7 (**Fig. 4F**). This resulted in an intermediate phenotype between full depletion and no depletion, indicating that both extrafollicular and GC derived antibody can feed back onto ongoing GCs to limit the success of B cell clones binding the same epitope (**Fig. 4G**). Altogether, our data indicate that antibodies produced by a B cell clone as it participates in both extrafollicular and GC responses can negatively affect the expansion of GC B cells of the same specificity.

### Antibody-mediated feedback in ongoing GCs is epitope-specific

Depletion of anti-Sa titers partially rescued the decay of *Ig*^Sa^ in the GC without detectably changing GC size, implying that the effect of antibody in this system is epitope-specific. To formally test this, we engineered a second BCR knock-in mouse harboring specificity to another, non-overlapping epitope on HA. Because it bound to site “Cb,” an epitope that does not overlap with the Sa epitope^33^, we chose clone 73.1 from our panel of 20 mAbs (**Fig. S3C**) for further characterization. Overlapping cryo-EM structures of Fabs 1.1 and 73.1 complexed with PR8 HA confirmed the footprints of the two antibodies did not overlap, and that clone 73.1 is Cb-specific (**Fig. 5A**). We therefore refer to clone 73.1 as “clone Cb,” To ensure that mAbs Sa and Cb did not inhibit each other’s binding due to steric hinderance, we performed competitive binding ELISAs. Even at 100-fold higher concentrations of competing mAb, Cb did not inhibit binding of Sa to HA and vice versa (**Fig. 5B**). Absence of competition between Sa and Cb mAbs was also verified by bio-layer interferometry, where binding of one antibody to HA did not prevent the binding of the other (**Fig. S5A**). The affinity of Fab Cb was approximately 30 nM (**Fig. 5C**), similar to the affinity of Fab Sa, increasing the likelihood that these two clones would simultaneously partake in a GC reaction^5^.

**Figure 5.**
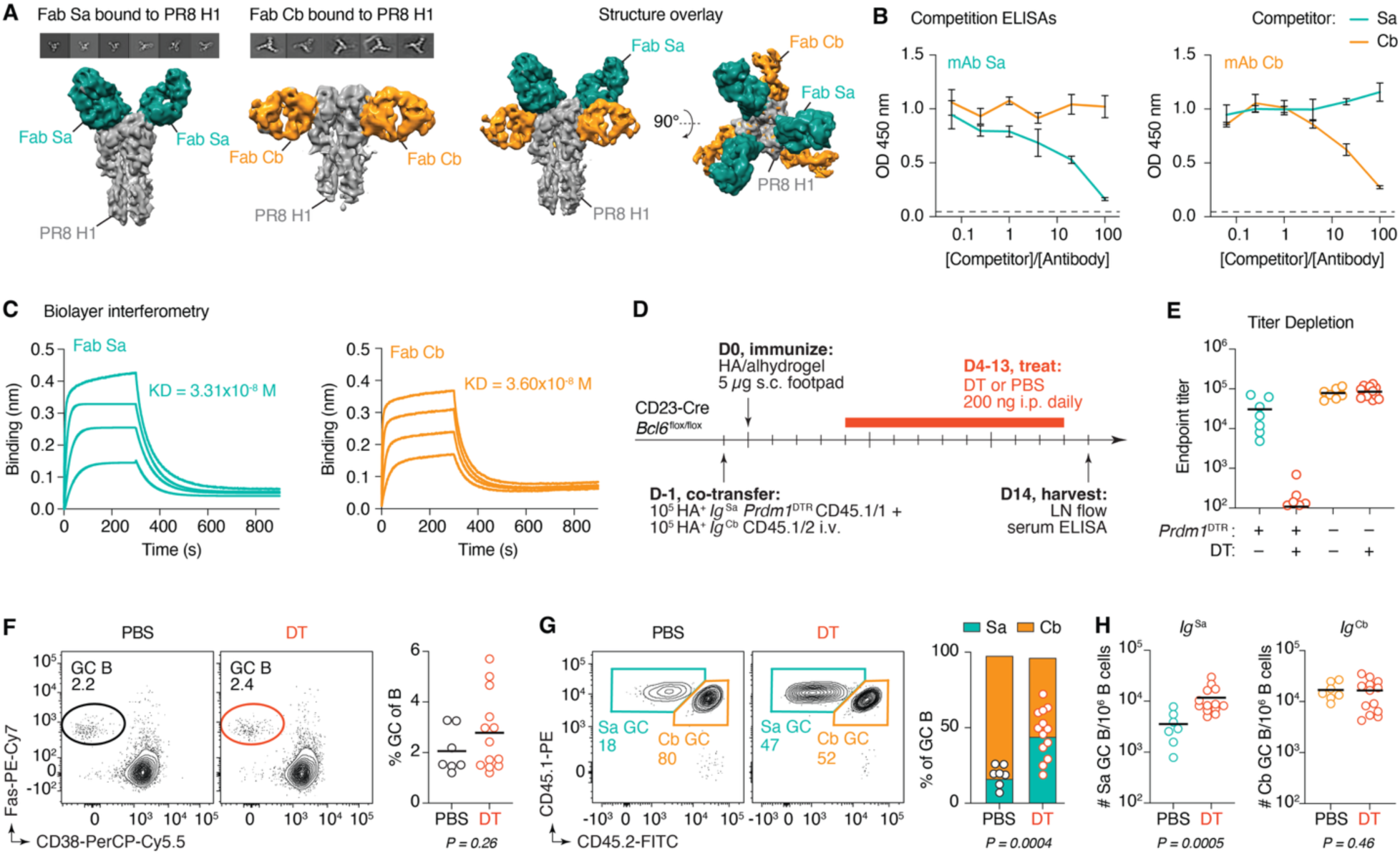
Antibody mediated suppression of competing GC responses is epitope-specific. **(A)** Representative 2D class averages showing different views of the PR8 H1 and Fab Sa or Fab Cb complex, along with 3D surface representations of complexes alone (*left*) and overlayed (*right*). **(B)** Competition for HA binding in ELISA between mAbs Sa and Cb. *Left*, mAb Sa is biotinylated; *right*, mAb Cb is biotinylated. Data are pooled from two independent experiments with technical duplicates. Lines connect means, and error bars represent SDs. **(C)** Biolayer interferometry of Fabs Sa and Cb binding to HA. Shown is one of two representative experiments. **(D)** Experimental setup for panels (E-H). **(E)** Endpoint titer dilutions of Sa and Cb IgG at 14 dpi in the presence or absence of DT treatment. **(F)** Representative flow cytometry plots (*left*) and quantification of GC size (*right*) at 14 dpi. **(G)** Representative flow cytometry plots (*left*) and quantification of *Ig*^Sa^ and *Ig*^Cb^ GC occupancy (*right*). **(H)** Number of *Ig*^Sa^ (*left*) and *Ig*^Cb^ (*right*) GC B cells per million LN B cells under steady state or upon depletion of *Ig*^Sa^ plasmablasts. (E-H) Each symbol represents one mouse. P-values are for Student’s T-test. (E-H) Results are pooled from 4 independent experiments, with n = 4 mice per group. Bars represent means. Mice with GC sizes less than 0.75 SD below the mean were excluded from analysis.

As we did for clone Sa (**Fig. 3A**), we engineered mice carrying the VDJH and VJκ sequences of clone Cb in their endogenous *Igh* and *Igk* loci, respectively. We crossed the resulting animals to the CD45.1 allelic marker to obtain *Igh*^Cb/+^.*Igk*^Cb/–^ CD45.1/2 (*Ig*^Cb^) mice, which we used as donors for adoptive transfer experiments. Soluble Sa mAbs did not compete with *Ig*^Cb^ B cells binding to HA tetramer via flow cytometry, and vice-versa (**Fig. S5B**). To test the epitope specific effects of antibody-mediated feedback, we set up an adoptive transfer system in which we could deplete only the antibodies secreted by *Ig*^Sa^ B cells in the presence of *Ig*^Cb^ competitors. We co-transferred 10^5^ *Ig*^Sa^*.Prdm1*^DTR^ HA^+^ B cells and 10^5^ *Ig*^Cb^ HA^+^ B cells into CD23-Cre; *Bcl6*^fl/fl^ hosts one day prior to footpad immunization with HA. We began dosing with DT at day 4 and analyzed popliteal GCs 2 weeks after immunization (**Fig. 5D**). Antibody depletion was specific to the clone harboring *Prdm1*^DTR^, as administration of DT eliminated anti-Sa titers without effecting anti-Cb titers (**Fig. 5E**). GC size was not detectably different in the absence of Sa antibody, although variability was too high to establish the absence of an increase (**Fig. 5F**). Despite their similar affinities for HA, *Ig*^Sa^ B cells were moderately outcompeted by *Ig*^Cb^ B cells, so that a 50:50 adoptive transfer of *Ig*^Sa^ and *Ig*^Cb^ B cells resulted in only 17% of *Ig*^Sa^ B cells in the GC at 14 dpi. However, depletion of anti-Sa titers resulted in a 2.7-fold expansion of clone *Ig*^Sa^, from 17% to 45% of GC B cells (**Fig. 5G**). Whereas the minor population of *Ig*^Sa^ GC B cells expanded approximately 3-fold in the absence of Sa antibody, the number of *Ig*^Cb^ GC B cells remained stable, indicating that secreted antibody specifically lowered the abundance of *Ig*^Sa^ GC B cells (**Fig. 5H**). We conclude that antibody-mediated suppression during an ongoing humoral response affects specifically B cells that compete with that antibody for antigen binding.

## DISCUSSION

We use a series of novel mouse genetics tools to investigate how antibodies generated by a contemporaneous immune response feed back onto their GCs of origin to regulate intraclonal and interclonal competition. Our data suggests that competition between B cells and antibody produced by PCs is not a primary driver of affinity maturation. However, it can influence the clonal composition of ongoing GCs by restricting access to antigen by B cells that recognize identical or overlapping epitopes.

We employed three strategies to investigate the effect of antibody-mediated feedback on affinity maturation. The first two quantify affinity-enhancing mutations at the *Ig* sequence level in GC B cells obtained from the functionally monoclonal response to NP-OVA immunization and from a truly monoclonal response by adoptively transferred HA-specific B cells. The third assessed the avidities of a *bona fide* polyclonal response to HA. In each of these experimental setups, we found no significant differences in the affinities or avidities of GC B cells when response-specific antibodies were depleted. Depletion of PCs after administration of DT usually resulted in reductions in antibody titers in the order of 100-fold or higher. Although depletion was not always complete, it consistently produced changes in the clonal composition of GCs (Figs. 4 and 5). We therefore consider it unlikely that residual antibody— whether derived from surviving PCs or secreted in small amounts by GC B cells themselves—would be sufficient to drive affinity maturation forward, unless the threshold for such an effect is substantially lower than that required to modulate interclonal competition. Moreover, our readouts are sufficiently sensitive to detect biologically meaningful changes in affinity, as demonstrated for both NP- and HA-specific responses^6,31,43^. Nevertheless, we cannot exclude the possibility that antibody-mediated feedback exerts a subtle influence on affinity maturation that falls below the detection threshold of our assays. However, such effects are unlikely to substantially impact the overall effectiveness of an immune response.

The reasons for the partial discrepancy between our findings on affinity maturation and the seminal work of Toellner et al.^22^ are not entirely clear. One possibility is that the effect of the exogenously injected high-affinity, high-avidity IgMs used in that study is more pronounced than that of the low-avidity IgGs that dominate endogenous responses in mice. This would be the case if the avidity advantage that B cells themselves have for binding antigens closely arrayed on the FDC network negates the effects of competing low-avidity IgG antibodies, even if these have higher nominal affinities than the B cell’s surface Ig. A recent study shows that 2-dimensional BCR-antigen on- and off-rates observed when antigen and surface Ig are both membrane bound cannot be reliably deduced from 3-dimensional antibody-antigen binding rates, highlighting the challenge in extrapolating the competitive dynamics between soluble antibodies and their membrane bound counterparts from conventional affinity measurements^44^.

In contrast to the rules of intraclonal competition—which we understand to be based on relative comparisons between the affinities of GC B cells that bind to the same epitope^6^—few if any rules governing interclonal competition in the GC have been defined thus far. Affinity alone does not explain the dominance of certain clones over others in the GC. B cell clones with lower nominal affinities frequently share GCs with much higher-affinity competitors, even when they recognize the same antigen^9,21,38,45,46^, implying that biochemically-measured affinity may not be a good proxy for GC competitiveness when different clones are compared. Our findings show that antibody-mediated feedback can influence these dynamics. Depletion of antibody from a polyclonal response induced by immunization with NP resulted in greater retention of the dominant Igλ^+^ clonotype. Similarly, preventing *Ig*^Sa^ cells from producing antibody increased the competitiveness of that clone in GCs when facing either a full polyclonal repertoire or a single monoclonal population specific for a distinct epitope. We suggest that, in the same way as pre-existing antibody— generated by either infusion of monoclonal antibodies or by a previous primary response—suppresses the entry of competing clones into nascent GCs^15–17,19,21^, antibodies formed during an ongoing B cell response can specifically mask the epitopes of the B cell clones that produced them, providing some degree of competitive advantage to clones specific for other, non-masked, epitopes. Such an effect, first predicted by quantitative modeling^39^, may act in concert with the reduction of affinity thresholds for GC entry by non-competing antibodies^18,45^ to create a negative feedback loop that attenuates the Darwinian push towards excessive clonal focusing, promoting clonal diversity in the antibody response.

## Supporting information

Supplemental Spreadsheet 1 - Ig sequences

## ACKNOWLEDGMENTS

We thank all members of the Victora laboratory as well as J. Chaudhuri (MSKCC), J.M. Rock (RU), J.V. Ravetch (RU), D. Mucida (RU), N. Wingreen (Princeton University), and A. Gupta (RU) for helpful discussion, M. Ebrahim, J. Sotiris, and H. Ng at the Rockefeller University Cryo-electron Microscopy Resource Center for assistance with structure data acquisition, the Rockefeller University Comparative Biosciences Center for mouse housing, and all Rockefeller University staff for their continuous support. We thank K. Gordon and J.-P. Truman for single-cell sorting, G. Kelsoe for NB-21.2D9 cells, J. Yewdell for recombinant Δ4 mutant HA proteins, J. Chaudhuri and M. Busslinger (IMP Vienna) for Cd23-cre mice, M. Reth (U. Freiburg) for *Cd79a*^Cre^ mice, C.-A. Reynaud and J.-C. Weill (U. Paris-Descartes) for *Aicda*^CreERT2^ mice, and A. Dent (Indiana U.) for *Bcl6*^flox^ mice. This study was funded by NIH/NIAID grants R01AI119006, R01AI173086, and R01AI180451 to G.D.V. Work in the Victora laboratory is additionally supported by the Robertson Foundation and the Stavros Niarchos Foundation (SNF) as part of its grant to the SNF Institute for Global Infectious Disease Research at The Rockefeller University. A.B. was supported by NIAID grant F30AI157448, Louis and Rachel Rudin Fellowship in Immunology, and NIGMS Medical Scientist Training Program grant T32GM007739. J.Bi. was supported by a Boehringer Ingelheim Fonds PhD fellowship and by the National Cancer Institute of the National Institutes of Health under Award Number F99CA305580. A.K.T. was supported by a scholarship from the German National Academic Foundation. G.D.V. is a Burroughs-Wellcome Investigator in the Pathogenesis of Infectious Disease and an HHMI Investigator.

## AUTHOR CONTRIBUTIONS

A.B. and J.Bi. performed most experimental work and data analysis, with help from T.L., A.K.T, A.H., J.-J. S., N.H.A. C.L.F., and essential input from R.V.H.C. and L.M. Nojima culture experiments were performed and analyzed by A.B. and T.L., with assistance from A.H. Data acquisition and analysis for CryoEM was performed by L.U. Transgenic mice used throughout this study generated in house were produced by J.Bo. A.B. and G.D.V. conceptualized the study; A.B., J.Bi. and G.D.V designed all experiments, interpreted the data, and wrote the manuscript. All authors reviewed and edited the final manuscript.

## MATERIALS AND METHODS

### Mice

Wild-type C57BL/6J (CD45.2) mice were purchased from the Jackson Laboratory (strain 000664). The following strains were kindly provided to us: *Aicda*^CreERT2^ by J.-C. Weill and C.-A. Reynaud^28^ (U. Paris-Descartes), CD23-Cre by M. Busslinger (IMP Vienna) and J. Chaudhuri (MSKCC)^26^, *Bcl6*^flox/flox^ by A. Dent^37^ (Indiana U.), and *Cd79a*^Cre^ by M. Reth (U. Freiburg)^36^. CD45.1 mice (*Ptprc*^K302E^ JAXBoy; Jackson Laboratory strain 033076) were bred and maintained in our laboratory. All mice were held under specific pathogen-free conditions at the immunocore clean facility at the Rockefeller University. Rockefeller University’s Institutional Animal Care and Use Committee approved all mouse procedures.

### Immunizations, Infections, and Treatments

Six to twelve-week-old male and female mice were subcutaneously immunized in the right footpad with 5 µg HA supplemented with 1/3 volume alhydrogel adjuvant (Invivogen) or 10 µg NP(16-19)-OVA (Biosearch technologies) supplemented with 1/3 volume alum adjuvant (G Biosciences). For NP-OVA pre-immunization, mice were immunized with 50 µg NP(16-19)-OVA supplemented with 1/3 volume alum adjuvant (G Biosciences) intraperitoneally. For IgY immunizations, mice were immunized subcutaneously in the hind footpads with 10 µg IgY (Exalpha Biologicals) supplemented with 1/3 volume alum adjuvant (G Biosciences). In *Aicda*^CreERT2^.*Prdm1*^LSL-DTR^ mice, the stop cassette in *Prdm1*^LSL-DTR^ was excised by oral gavage of 200 µl tamoxifen (Sigma) dissolved in corn oil at 50 mg/ml, on days 2, 5, 8, and 11 post-immunization. Diphtheria toxin (Sigma) was resuspended in PBS at 1 mg/ml, and mice were dosed daily beginning at day 4 with 200 ng intraperitoneally. Blood samples for ELISA quantification of titers were collected by cheek puncture into microtubes prepared with clotting activator serum gel (Sarstedt). For influenza infections, mice were anesthetized with ketamine/xylazine diluted in sterile PBS 1X (Gibco, Inc.) and infected intranasally with 33 PFU of mouse-adapted influenza PR8 virus (provided by M. Carroll, Harvard Medical School).

For adoptive B cell transfer, spleens of naive *Ig*^Sa^ and *Ig*^Cb^ mice were harvested and mechanically homogenized through a 70-um strainer. Red blood cells were lysed with ACK buffer (Thermo Scientific). B cells were isolated by negative selection with anti-CD43 beads (Miltenyi Biotec). 100 µl PBS containing 10^5^ HA tetramer-binding B cells or 5x10^5^ IgY-binding B cells from *Ig*^chIgY^ knock-in mice was injected intravenously into recipient hosts, one day prior to immunization or infection.

### Mouse engineering

*Prdm1*^LSL-DTR^, *Ig*^Sa^, *Ig*^Cb^, and *Igk^−^* mice were generated in our laboratory. For *Prdm1*^LSL-DTR^, we designed the allele as indicated in Fig. 1A and Fig. S1A, as described in the text. The 1392-nucleotide double-stranded DNA template (including 5’ and 3’ homology arms, each 200 nucleotides long) and the CRISPR guide-RNA (TGAAAATCTTAAGGATCCAT) were purchased from IDT. Single-stranded DNA was prepared from the double-stranded *Prdm1*^LSL-DTR^ template (Fig. S1) using the Guide-it Long ssDNA Production System v1 (Takara) as per the manufacturer’s instructions. Single-stranded DNA template and the CRISPR guide-RNA were prepared according to the Easi-CRISPR gene targeting method^34^ and injected into C57BL/6J zygotes. Resulting mice were backcrossed for at least three generations onto C57BL/6J before being used in final experiments. The constitutive *Prdm1*^DTR^ allele was generated by crossing *Prdm1*^LSL-DTR^ to *Cd79a*^Cre^, which leads to occasional deletion of the stop cassette in the germline. For generation of B cell receptor knock-in *Ig*^Sa^ and *Ig*^Cb^, we followed a similar protocol but this time used two CRISPR guides for each insertion in order to delete the J-segments of the native *Igh and Igk* loci. The guides use for *Igh* were TCTCTACTTCCTCATAGCTC and GGAGCCGGCTGAGAGAAGTT, and the guides used for *Igk* were CTGTGGTGGACGTTCGGTGG and AAGACACAGGTTTTCATGTT. *Ig*^Sa^ and *Ig*^Cb^ *Igh* and *Igk* template constructs are shown in Fig. 3A. To produce the *Igk^-–^* allele, we injected C57BL6/J zygotes with an sgRNA targeting *Igkc* (GTTCAAGAAGCACACGACTG), and screened mice for indels causing a frameshift in the *Igkc* locus^47^.

### Flow Cytometry and Cell Sorting

Cell suspensions from LNs were obtained through mechanical disruption with micropestles (Axygen). Cells were resuspended in PBS supplemented with 0.5% BSA and 1 mM EDTA and first incubated with Fc-block (rat anti-mouse CD16/32, clone 2.4G2, Bio X Cell) for 30 min on ice. After washing with PBS, cells were resuspended in a mix containing various fluorescently labeled antibodies for 30-60 min on ice. Cells were filtered e and washed with the same buffer before analysis on a BD FACS Symphony A5 cytometer or single cell sorted using a BD FACS Symphony S6. Data were analyzed using FlowJo v.10 software.

### Generation of Recombinant Proteins and mAbs

A CHO cell protein expression system was used to generate cysteine-crosslinked^48^ recombinant HA for immunization, as described in detail in a previous publication^49^. The same procedure was followed to generate SaΔ4 and CbΔ4 HA variants. Thrombin cleavage was used to remove C-terminal domains not native to HA (foldon, Avi-tag, His-tag) for use in immunization. HAs were then purified via FPLC, flash frozen in PBS using liquid nitrogen, and stored at –80°C. HA was used for ELISA was not thrombin treated. To create HA tetramers for use in flow cytometry, a non-cysteine-stabilized version of PR8 HA harboring the Y98F mutation that abrogates binding to sialic acid was produced as above. This HA was biotinylated at its Avitag using BirA-500 ligase (Avidity) and desalted by Zeba column purification (Thermo Fisher). Biotinylated HA was incubated with Streptavidin-APC in PRC for 30 min at room temperature at a molar ratio of 4 HA trimers to 1 Strepavidin APC. The plasmid used for HA cloning and expression (pVRC8400) was kindly provided by A. McDermott (VRC/NIAID/NIH). mAb heavy and light chain sequences were synthesized and assembled into Ig production plasmids by Twist Biosciences. 293F cells were transfected with plasmids, and mAbs and Fabs (his-tagged) were purified using protein-G or Ni-NTA (both from Cytiva) affinity chromatography, respectively, as described previously^9^.

### Antibody Binding Measurements

Bio-layer interferometry (BLI) was performed on an Octet RED96 instrument (ForteBio) to determine affinities of Fabs bound to influenza HA. Briefly, single site-biotinylated HA was loaded onto High Precision Streptavidin (SAX) Biosensors (ForteBio) until binding reached 1 nm. Association was measured by submerging sensors into wells containing various concentrations of Fabs (160 nM – 40 nM in PBS 0.1%BSA/0.02%Tween 20) for 300 s. Disassociation was measured by transferring sensors into empty buffer solution for 600 s. KD values were determined using the global fit 1:1 binding algorithm provided with the Octet data analysis software. For epitope binning experiments, HA-loaded sensors were submerged into 160 nM of one antibody and then immediately submerged in buffer solution containing 160 nM of the other antibody without an intervening disassociation step.

### Cryo-EM Sample preparation, data collection and processing

A volume of 3 µL of 3.3 µM HA mixed with 10 µM Fab in PBS was applied to plasma-treated Quantifoil R2/2 300-square-mesh copper grids. The grids were blotted for 3 to 5 seconds at 100% humidity and 4°C using a Vitrobot IV (FEI) before being plunged into liquid ethane.

A total of approximately 1,000 micrographs of each complex were recorded using an FEI Arctica equipped with a Gatan K2 Summit detector, utilizing Serial EM for automated data collection software (Mastronarde, 2005). Motion correction and dose weighting were performed using MotionCor2^50^. The dose- weighted summed micrographs were then used to estimate the contrast transfer function (CTF) parameters with CTFFIND4^51^. Subsequent data processing was carried out using RELION v3.0^52^.

Initially, a small subset of particles was manually picked and subjected to 2D classification. The resulting 2D references were used for autopicking all the micrographs. The autopicked particles underwent a cleaning process, which included multiple rounds of reference-free 2D classification to select the best particles. The cleaned particles were used to generate initial models, which were low-pass filtered to a resolution of 50 Å and used as a reference for 3D refinements. 3D refinement yielded a density map of Sa- and Cb-bound HA at resolutions of 6.7 Å and 5.5 Å, respectively. For the HA protein and Fabs, PDB 1RU7 and PDB 4HLZ respectively were rigid-body fitted into refined density maps within Chimera^53^.

### Single-GC B cell “Nojima” cultures

The NB-21.2D9 cell line expressing BAFF, CD40L, and IL-21 (kindly provided by G. Kelsoe, Duke University) was cultured in DMEM containing 10% heat-inactivated FBS and penicillin streptomycin solution (Corning). The day before sorting, feeder cells were detached, resuspended in OptiMEM, irradiated with 20 Gy and seeded into 96-well plates at 3,000 cells per well in OptiMEM supplemented with 10% heat-inactivated FBS, 2 mM L-glutamine, 1 mM sodium pyruvate, 50 uM 2-ME, penicillin streptomycin solution, 10 mM HEPES, MEM vitamin solution (Sigma) and EMM non-essential amino acids (Gibco). After single- cell sorting, 150 µl of supplemented OptiMEM with 30 µg/ml LPS (Sigma-Aldrich, #L6511) and 4 ng/ml IL- 4 (Fisher Scientific, #404-ML) was added to each well. Supernatants were harvested 7 days after sorting and screened for IgG1 and HA reactivity by ELISA, as described below. For the experiment shown in Figure 3K, the yield was 357 IgG1^+^ cultures out of 2,134 sorted cells.

### ELISA

To determine antibody levels from supernatants of single GC B cell cultures, 96-well high-binding half-area microplates (Greiner) were coated overnight at 4°C with antigen of capture antibody in PBS (25 µl per well). After overnight incubation, plates were blocked with 2.5% BSA (Sigma) in PBS for 2 hours at room temperature. For single GC B cell culture experiments, 25 µl of undiluted supernatant was added to wells coated with 1 µg/ml goat anti-mouse Ig (Southern Biotech) or HA. After washing three times with PBS + 0.05% Tween20, plates were incubated at room temperature for one hour with detection antibody goat anti-mouse IgG1-HRP (Southern Biotech). Following two washes with PBS + 0.05% Tween20 and another two washes with PBS, plates were developed with slow kinetic form TMB (Sigma). Absorbance at 450 nm was measured on a Fisher Scientific accuSkan FC plate reader after the reaction was halted with 1N hydrochloric acid. To determine antibody levels in the serum of mice, antigen coated ELISA plates were incubated with 3-fold serial dilutions of serum samples starting at 1/100. For total IgG1 quantifications, serum dilutions were incubated onto Ig capture coated ELISA plates. Standard curves of mouse IgG1 were created by incubating Ig capture coated ELISA plates with two-fold dilutions of unlabeled mouse IgG1(Southern Biotech), beginning at a concentration of 2 µg/ml. To perform competitive ELISAs, mAbs Sa and Cb were biotinylated at a 5:1 biotin:mAb molar coupling ratio as per the manufacturer’s instructions (EZ-link Sulfo NHS-LC-LC-Biotin, Thermo Fisher). 1 nM of biotinylated antibodies and various concentrations of non-biotinylated antibodies were co-incubated for 30 min at room temperature before being pipetted into HA coated ELISA plates. Detection of bound biotinylated antibody was performed with Streptavidin-HRP (Biorad) at a 1:20,000 dilution.

### Single-cell *Ig* sequencing

Index-sorting of single GC B cells was performed into 96-well plates containing 5 ml TCL buffer (QIAGEN) supplemented with 1% β-mercaptoethanol. Nucleic acids were extracted using SPRI beads as previously described. RT maxima reverse transcriptase (Thermo Scientific) with oligo(dT) as a primer was used for reverse-transcription into cDNA. A forward primer mix comprising of consensus sequences for all V-regions and reverse primers for each isotype was used to PCR amplify *Igh* genes. To generate sequences for antibody production, *Igk* genes were amplified separately where needed. Afterwards, 5-nucleotide barcodes were introduced by PCR to label Ig-sequences with plate- and well-specific barcodes. The forward and reverse primers contained barcodes to identify the plate and row number, and barcodes to identify the column position, respectively (adapted from ^54^). A final PCR step was performed to incorporate Illumina paired-end sequencing adapters into single-well amplicons. PCR-products from all plates were pooled together and subsequently cleaned-up using a 0.7x volume ratio of SPRI beads. Sequencing was performed on the Illumina Miseq platform with a 500-cycle Reagent Nano kit v2 as per the manufacturer’s instructions. Primer sequences are provided separately.

### Analysis of sequencing data

PandaSeq was used to assemble paired-end sequences and processing was performed with the FASTX toolkit. Barcode sequences were used to assign the resulting demultiplexed and collapsed reads to their respective plates and wells. High-count sequences were analyzed for every single cell/well. To determine the V(D)J arrangements and the number of somatic mutations compared to putative germline precursors, Ig heavy chain and Ig light sequences were aligned to the online databases (IMGT; Vbase2). Sequences that shared VH/JH genes, had the same CDR3 lengths, and contained at least 75% CDR3 nucleotide identity were grouped and classified automatically into clonal lineages. Manual curation was performed based on features including V-region SHM patterns and stretches of mismatches at the junctional regions, resulted in further joining of sequences deemed to belong to the same clone, but which fell below 75% CDR3 nucleotide identity. Only cells with productively rearranged *Igh* genes were used for VH mutation analyses. To detect individual mutations, sequences were translated and aligned to the original unmutated VDJ sequence.

### Data analysis

Figure legends indicate which statistical tests were used to compare conditions. Parametric tests were performed after assessing whether the data was normally or lognormally distributed, using Shapiro-Wilk tests. In the case of lognormal distributions (e.g., ELISA titers) the log-transformed values were used for calculating significance. Statistical analysis was carried out using GraphPad Prism v.10. Flow cytometry analysis was carried out using FlowJo v.10 software. Graphs were plotted using Prism v.10 and edited for appearance using Adobe Illustrator.

## SUPPLEMENTAL FIGURES AND LEGENDS

**Figure S1.**
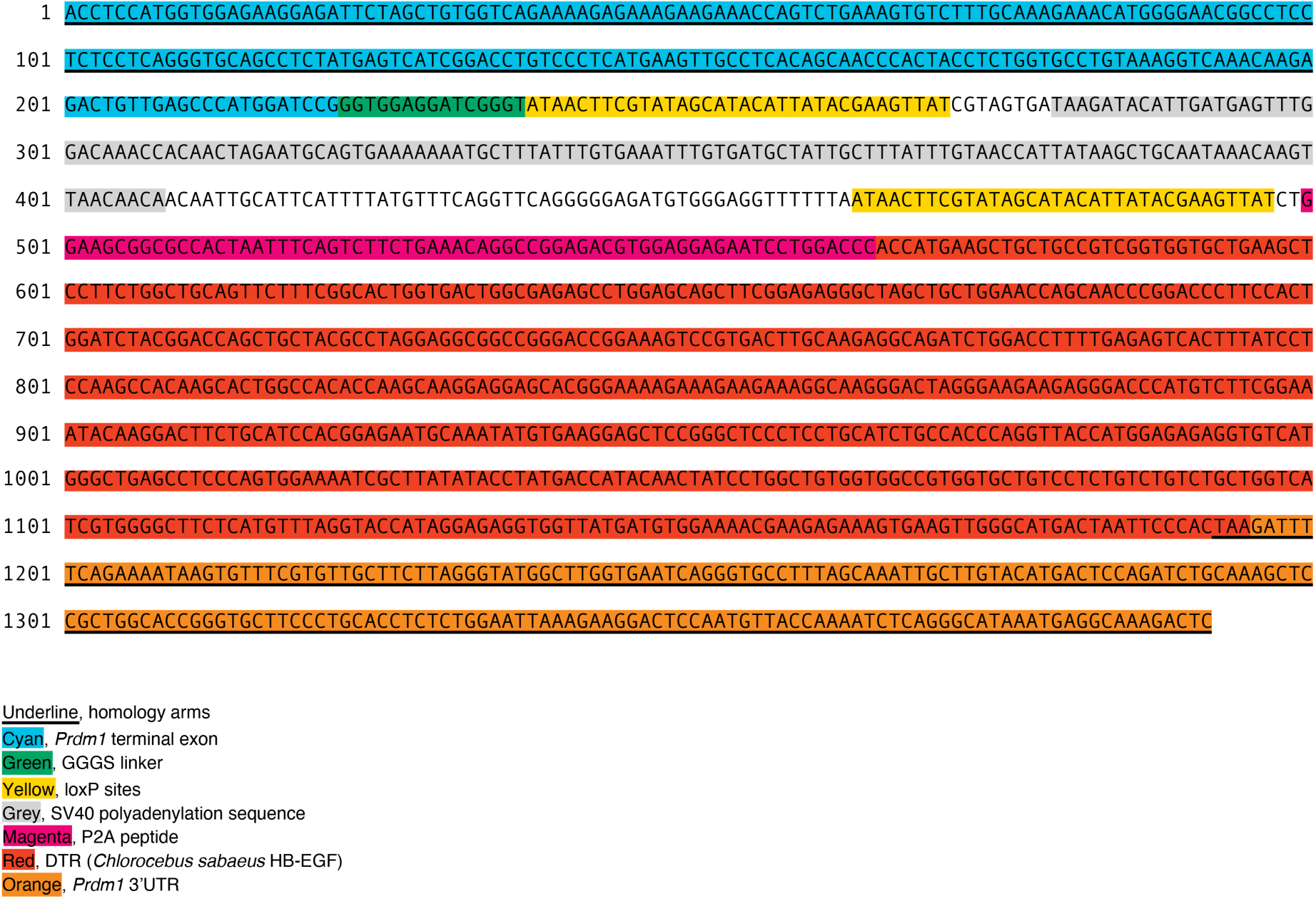
Annotated sequence for the repair oligo used to build the *Prdm1*^LSL-DTR^ allele.

**Figure S2.**
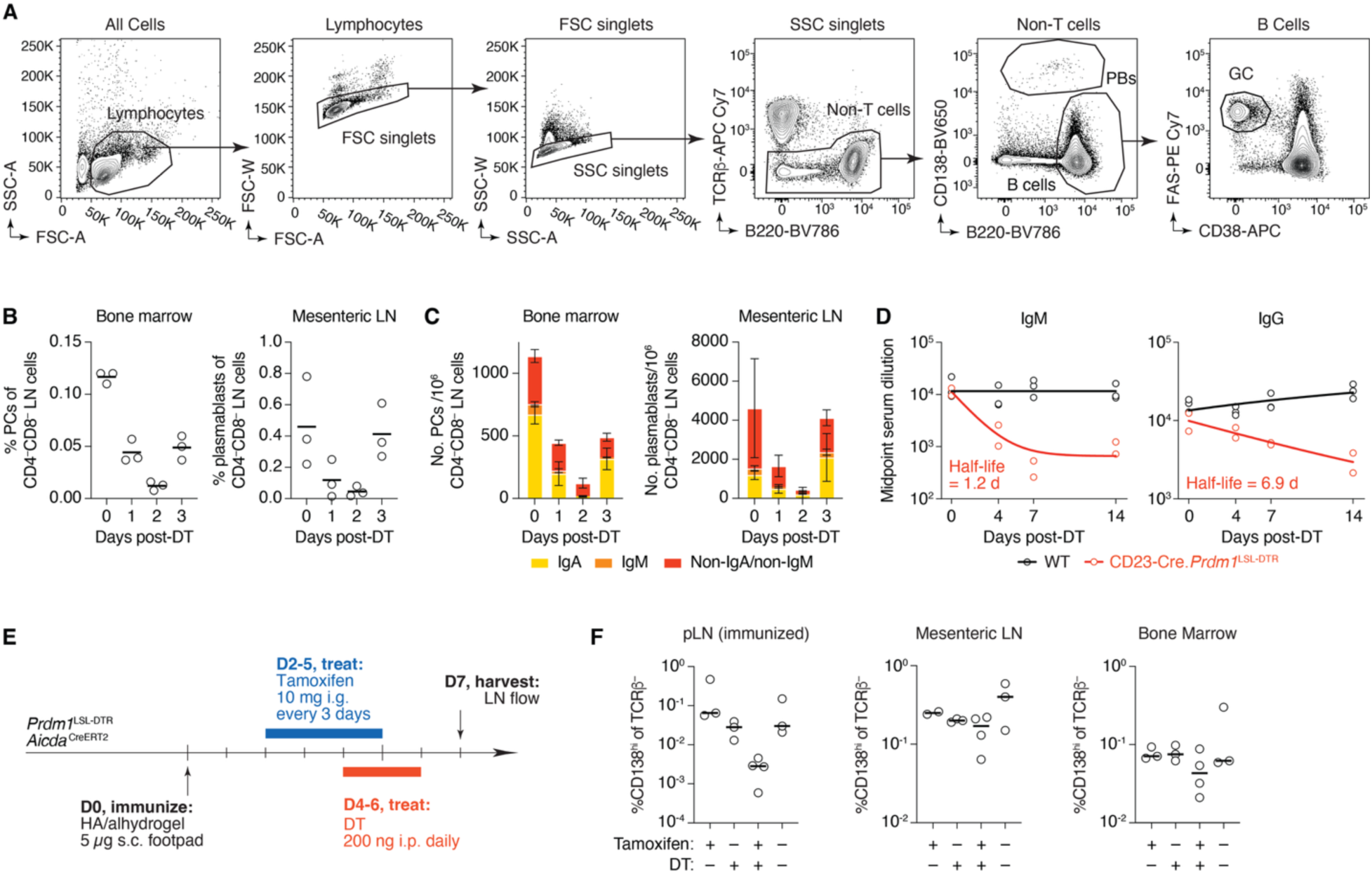
Characterization of plasma cell and antibody depletion in *Prdm1*^LSL-DTR^ mice. **(A)** Gating strategy for identification of LN plasmablasts and GC B cells. In some experiments, GC B cells were identified as CD38^low^ GL-7^+^, and in some experiments PCs were identified as CD4/CD8^−^CD138^hi^. **(B)** Kinetics of depletion and resurgence of bone marrow PCs and mesenteric LN plasmablasts following i.p. administration of 200 ng of DT to CD23-Cre.*Prdm1*^LSL-DTR^ mice. Each symbol represents one mouse. **(C)** As in (B), but following the counts per million LN cells of PCs and plasmablasts of each isotype. Non-IgA/non-IgM cells are presumably IgG^+^, given that the latter isotype is downregulated upon B cell differentiation to the plasmablast and PC fate. Error bars are S.D. for three mice as in (B). **(D)** Cre.*Prdm1*^LSL-DTR^ mice were given 200 ng of DT daily for the duration of the experiment and serum samples were collected at 0, 4, 7, and 14 days post-treatment. Midpoint serum titers of total IgM and IgG were determined by ELISA and the half-life of each isotype was determined using one-phase exponential decay. Each symbol represents one mouse, the line shows the fitted exponential curve. (**E**) Experimental setup. (**F**) Quantification of LN plasmablasts (*left and middle*) and bone marrow PCs (*right*) 7 days post HA immunization depending on treatment regimens as indicated below graphs. Each symbol represents one mouse. Results are pooled from two independent experiments with n=1-2 mice per group. Bars represent medians.

**Figure S3.**
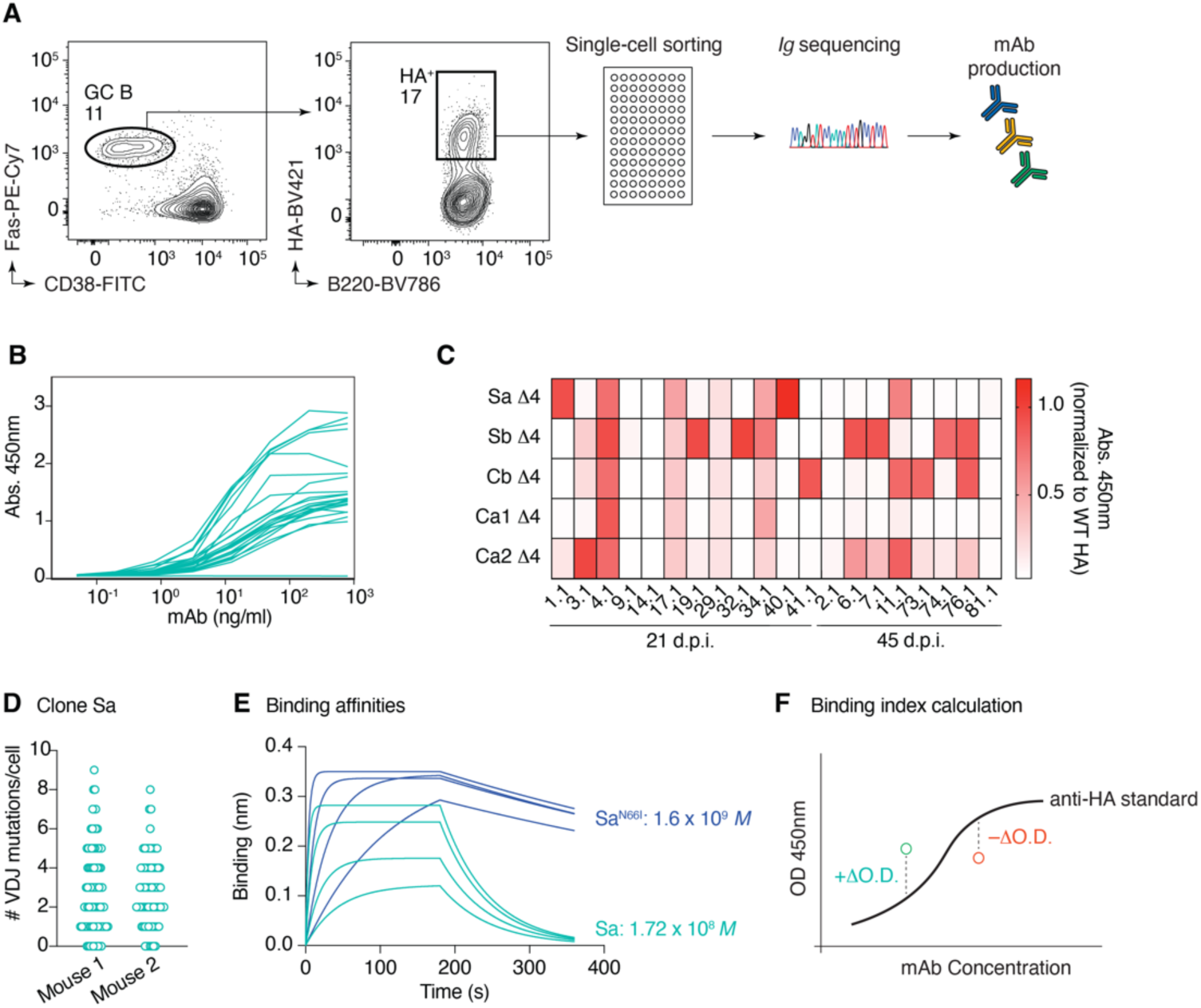
Characterization of monoclonal antibodies specific for influenza HA. **(A)** C57BL/6 mice were infected with PR8 influenza virus and mediastinal GC B cells were assayed by flow cytometry 21 and 45 days later. HA tetramer binding B cells were single cell sorted, their *Ighv* and *Igkv* loci were sequenced, and monoclonal antibodies were produced from members of 20 expanded clones. **(B)** Titration curves of candidate mAbs binding to recombinant PR8 HA by ELISA. **(C)** ELISA of mAbs binding to five Δ4 mutants, in which all but one of the five antigenic sites of PR8 HA are mutated. mAbs were assayed at a concentration of 1 μg/ml. Sequences for (A) and (C) are available in Supplemental Spreadsheet 1. **(D)** Number of *Igh* nucleotide mutations per cell in *Ig*^Sa^ GC B cells 18 dpi with HA. **(E)** Binding kinetics curves of Sa sequence and N66I mutation Fabs as measured by bio-layer interferometry. Fabs were assayed at concentrations of 160, 120, 80 and 40 nM. **(F)** Schematic demonstrating calculation of binding index. Black curve represents binding of reference mAb Cb to HA in ELISA. mAbs with better binding than Cb will be higher than the curve (green circle), while mAbs with decreased binding will be lower (red circle).

**Figure S4.**
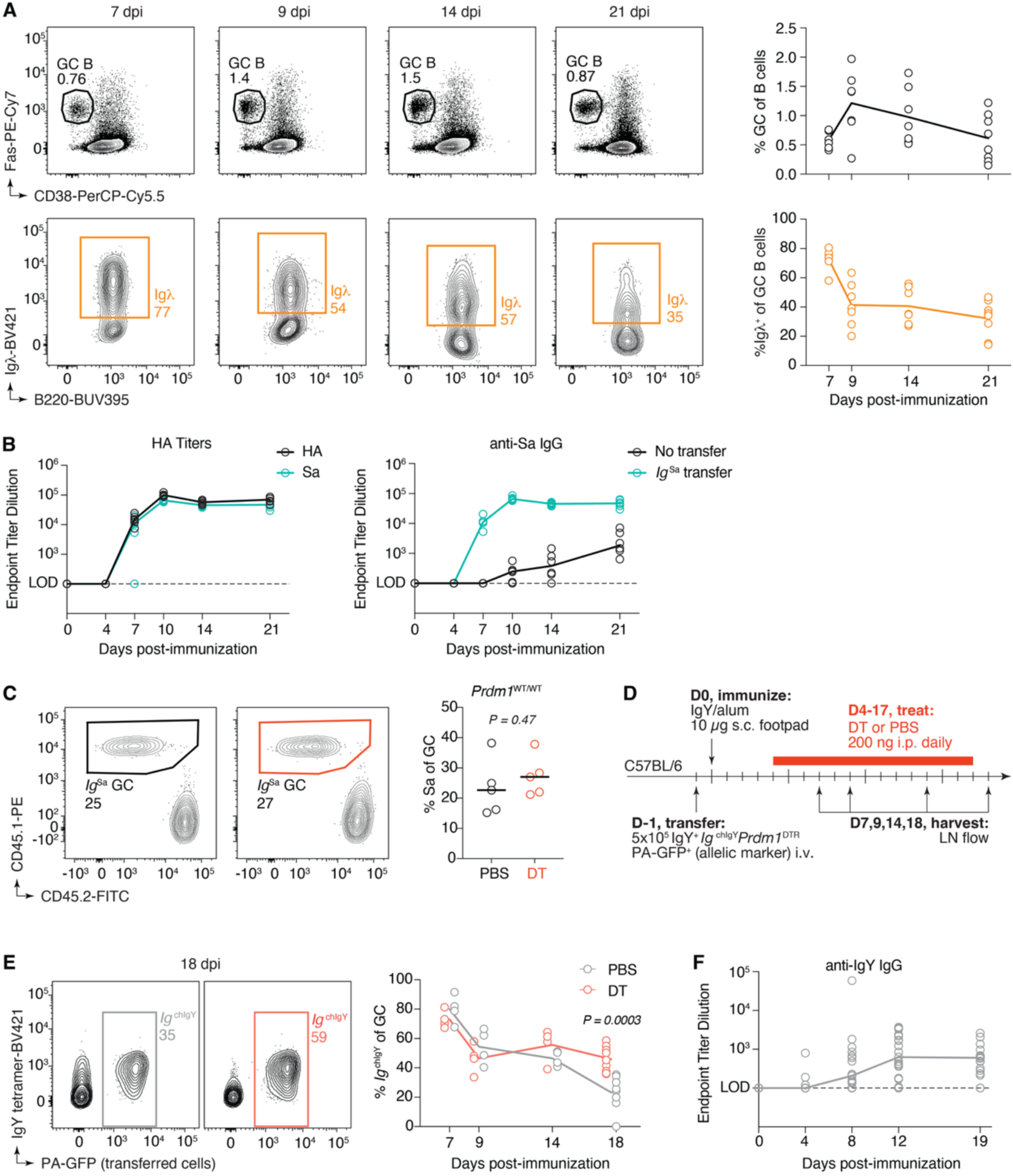
Antibody mediated feedback is not limited to the Sa epitope. (A) Immunization time course with NP- OVA following Igλ^+^ GC B cells in C57BL/6 mice. Top row shows representative flow cytometry plots and quantification of GC size and bottom row shows representative flow cytometry plots and quantification of Igλ^+^ GC occupancy. (B) Sa-specific and total HA IgG1 titers in C57BL/6 mice following *Ig*^Sa^ transfer and HA immunization. (C) Same experimental setup as in 4D, but *Ig*^Sa^ cells do not carry the *Prdm1*^DTR^ allele. Shown are representative flow cytometry plots (*left*) and a quantification (*right*) of *Ig*^Sa^ GC B cells 18 dpi. (D) Experimental setup for panel (E). (E) Representative flow cytometry plots and fraction of *Ig*^chIgY^ GC B cells at 7,9,14 and 18 dpi with IgY. PAGFP-transgenic mice are used as a marker for flow cytometry, without photoactivation. (F) IgY-specific IgG1 titers in C57BL/6 mice following *Ig*^chIgY^ transfer and IgY immunization. (A,D) Results are pooled from 2 independent experiments, with n = 2-4 mice per group. (B) Data are pooled from two independent experiments, with n = 3 mice per group. (C) Data are pooled from two independent experiments, with n = 2-3 mice per group. (F) Data are pooled from two independent experiments, with n = 8-11 mice per group. (B,F) Lines connect medians. In all plots, each symbol represents one mouse. P-values are for Student’s T-test.

**Figure S5.**
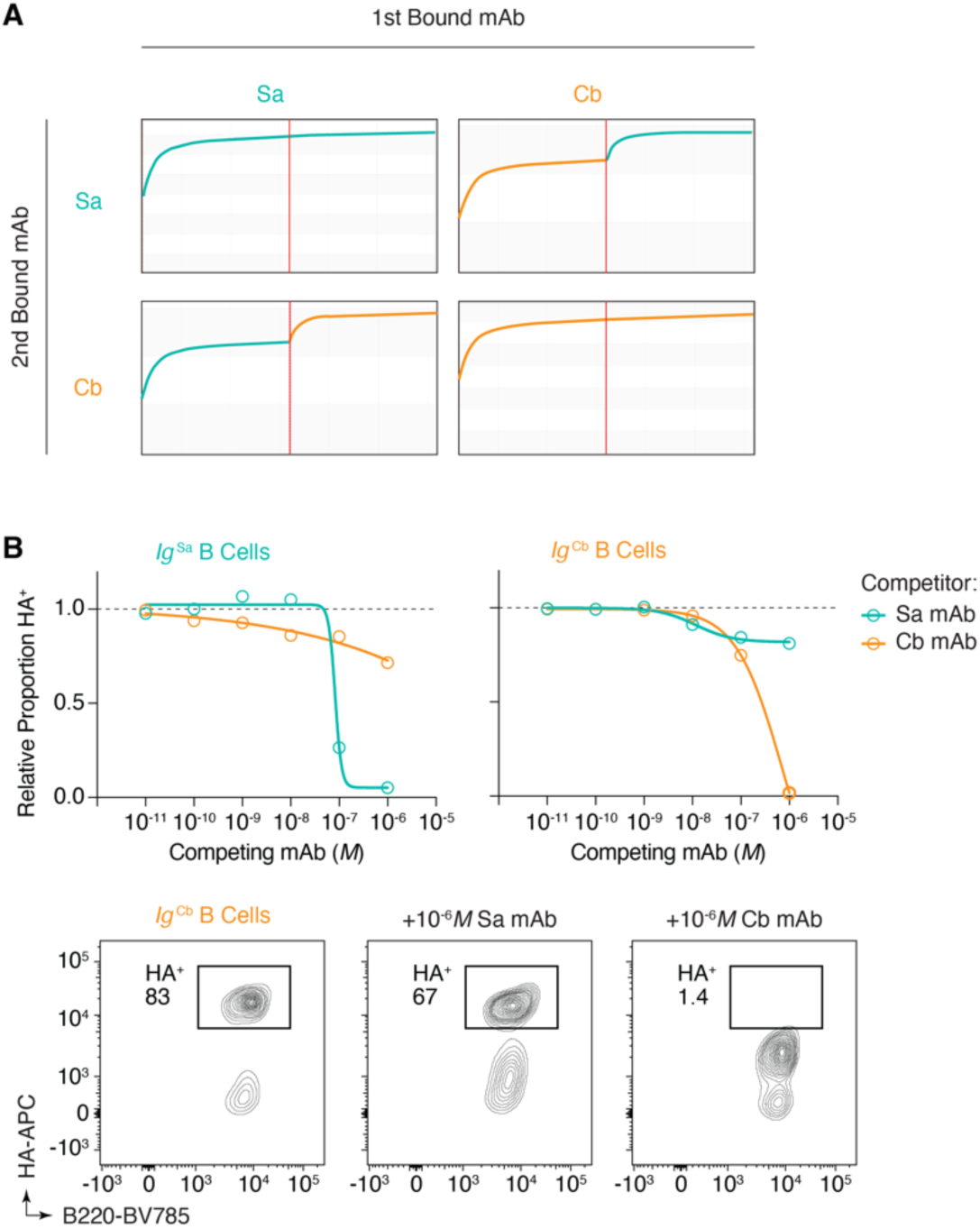
Characterization of two HA-specific monoclonal antibodies with non-overlapping epitopes. **(A)** Epitope binding via bio-layer interferometry. Binding of one mAb to PR8 HA was followed by binding with another, and binding to distinct sites is indicated by continued mass accumulation onto the sensor. **(B)** Flow-cytometry competition assays between *Ig*^Sa^ or *Ig*^Cb^ B cells and soluble mAbs for binding to HA tetramer. mAbs were co-incubated with tetramer at increasing concentrations, and inhibition of B cell staining was assessed. Proportion of HA^+^ is normalized to binding of HA in the absence of competing antibody.

